# Characterization of severe fever with thrombocytopenia syndrome virus Japanese isolate YG1 strain quasispecies using reverse genetics approaches

**DOI:** 10.1101/2024.02.01.578508

**Authors:** Sithumini M.W. Lokupathirage, Devinda S. Muthusinghe, Rakiiya S. Sarii, Olusola A. Akanbi, Kenta Shimizu, Yoshimi Tsuda, Kumiko Yoshimatsu

## Abstract

Three amino acid mutations have been identified in the isolated YG1 strain of severe fever thrombocytopenia syndrome virus (SFTSV), Gn (Y328H) accounts for 26.9% of the virus in patients’ blood, Gc (R624W) and L (N1891K) those are minor. To investigate viral properties caused by each mutation, we rescued viruses with one–three mutations. Mutations Y328H and R624W in GP increased the cell fusion activity and plaque size. Theses enhancement was more pronounced for both Y328H and R624W. The pseudotyped vesicular stomatitis virus coated with the SFTSV GP Y328H mutant showed lower infectivity in Vero E6 cells, which was compensated for by the additional R624W mutation. In the process of adaptation for virus with Y328H, the R624W mutation may be acquired. Moreover, only the viruses with the N1891K mutation in L showed significant CPE and the CPE was inhibited by the pan-caspase inhibitor, suggesting that caspase-dependent cell death occurred. Programmed cell death associated molecules caspase-1 and caspase-3 were induced in both CPE inducing and wild-type virus-infected cells. Furthermore, infection with the wild-type virus suppressed actinomycin D-induced cell death. These results suggest that SFTSV infected cells initiate programmed cell death, whereas wildt-ype virus may inhibit cell death. Furthermore, the N1891K mutation in L virus was outcompeted by a 10-fold less wild-type virus in Vero E6 cells indicating that it was not advantageous for viral survival in Vero E6 cells. Thus the quasispecies composition of SFTSV appeared to be influenced by propagative environment.

**Importance:** This study shows information on viral pathogenesis by analyzing quasispecies derived from one fatal case of severe fever with thrombocytopenia syndrome virus (SFTSV) infection. Observation with recombinant SFTSV altered Gn and Gc suggests that combining mutations may increase the viability of mutant viruses, selecting viruses to create a suitable population for propagation. The N1891K mutation in L protein of SFTSV was related to CPE appearance. On the other hand, wild-type virus which is major population in patient infection was suppressive for cell death. It was suggested that SFTSV has a mechanism to escape cell death for the prolonged viral propagation in infected cells. Although the mechanism is still unknown, it has been suggested that RNA virus polymerase might be involved in the regulation of cell death. This study proposed the mechanism underlying the adaptation to the environment and survival of virus as quasispecies.

## Introduction

Severe fever with thrombocytopenia syndrome (SFTS) is caused by the SFTS virus (SFTSV), a member of the order *Bunyavirales* and family *Phenuiviridae*. The SFTS is widely distributed in East Asian countries, including China, Japan, South Korea, and Vietnam (1–4). The mortality rate of SFTS is as high as 30% in Japan, and it is a public health concern in other countries (5–8). The SFTSV YG1 strain was isolated from the first patient with SFTSV in Yamaguchi Prefecture, Japan (2).

The SFTSV consists of three negative-sense segments, designated small (S), medium (M), and large (L). The S segment is an ambisense RNA encoding a nucleocapsid protein and a nonstructural protein (9); the M segment encodes two envelope glycoproteins (GP), Gn and Gc, whereas the L segment encodes an L protein that is an RNA-dependent RNA polymerase (RdRp).

In viruses of the family *Phenuiviridae*, entry into host cells is initiated by binding GPs (Gn and Gc) to cell surface receptors, and endocytosis occurs in a receptor-mediated manner (10–13). Following endocytosis, viral GPs fuse with the endosomal membrane under acidic conditions, causing low pH-dependent cell fusion in infected cells (14–16). Structural models of SFTSV Gn and Gc show that some amino acid residues are crucial for low pH-dependent cell fusion and syncytial formation (17, 18).

Following the cell entry, the infected cells of the Phenuiviruses, such as SFTSV and the Rift Valley fever virus, can cause cytopathic effects (CPE) (19, 20). Gao *et al.* reported that the cell death induced in human microglial HMC3 cells by infection of SFTSV was performed by the NOD-like receptor protein 3 inflammasome activation and leading to the secretion of interleukin (IL)-1β and pyroptosis (21). In contrast, most cell lines and primary cultures were not killed by SFTSV infection. Suzuki *et al.* reported that B cells are target cells in lethal SFTSV infections (22). However, transformed B cells and the primary culture of peripheral B cells did not die from SFTSV infection. Vero E6 and Huh7 cells used for virus isolation also did not show CPE, and the relationship between SFTSV infection and cell death remains largely unknown.

In a previous study, we established three subclones (E3, A4, and B7) from the YG1 strain using the limiting dilution method with pH-dependent cell fusion activity and CPE. Subclone E3 had a genome identical to that of the parental YG1 strain, whereas subclones A4 and B7 had two amino acid mutations in glycoproteins Gn (Y328H) and Gc (R624W).

Subclone B7 had another amino acid mutation, N1891K, in its L protein (19). Subclone A4 (Gn: Y328H and Gc: R624W) exhibited intense cell fusion activity under acidic conditions. In contrast, B7 (Gn: Y328H, Gc: R624W, and RdRp: N1891K) showed robust CPE, indicating the involvement of L protein N1891K in the difference in CPE between the subclones (19). In addition, the polarization of the amino acid at position 1891 of L protein is critical for its function, especially its polymerase activity (23). The Gn: Y328H mutation was found in 26.9% of the viral population in patient blood by next-generation sequencing (2, 24). However, after isolating the YG1 strain, the Gn: Y328H mutation rate reduced to approximately 10%. In contrast, two mutations, Gc: R624W and L: N1891K, appeared infrequently in the patient’s blood and the isolated YG1 strain. Furthermore, the roles of these mutations in the virological characteristics and population structure of SFTSV need to be clarified. Information on the emergence and survival of viruses whose pathogenicity changes due to mutations is essential for understanding their pathogenicity.

In this study, we developed recombinant viruses bearing single mutations using a reverse genetics system (9) to compare the unique virological characteristics of each mutation. In addition, the effects on quasispecies were evaluated, and how mutations are related to their impact on quasispecies is discussed.

## Materials & Methods

### 1. Cells and viruses

Vero E6 cells (ATCC C1008) were maintained in Eagle’s minimum essential medium (Gibco; Thermo Fisher Scientific, MA, USA) supplemented with 5% heat-inactivated fetal bovine serum (FBS) (Biowest, Nuaillé, France), 1% MEM non-essential amino acids (Gibco), 1% insulin-transferrin-selenium (Gibco; Thermo Fisher Scientific), 1% penicillin (50 units/mL), streptomycin (50 μg/mL; Sigma-Aldrich Co., St Louis, MO, USA), and 1% gentamicin (100 μg/mL; Sigma-Aldrich). BSR-T7/5 cells stably expressing T7 RNA polymerase were kindly provided by Dr. K. K. Conzelmann (Max-von-Pettenkofer Institute, Munich, Germany) (25). The cells were maintained in low-glucose Dulbecco’s modified Eagle medium (Sigma-Aldrich) supplemented with 10% FBS, 10% Tryptose Phosphate Broth (Gibco), and antibiotics (1 mg/mL geneticin (G418) (Nacalai Tesque, Kyoto, Japan) or 1% penicillin and streptomycin). Huh7 human hepatoma and 293T cells (Riken, Japan) were grown and maintained in Dulbecco’s modified Eagle medium (Thermo Fisher Scientific) supplemented with 10% FBS and 1% penicillin-streptomycin. All cells were cultured in a 5% CO_2_ incubator at 37 °C. The subclones of the YG1 strain, A4, E3, and B7, were used as controls. The viral infection experiments were performed in a biosafety level 3 (BSL-3) facility at the Institute for Genetic Medicine, Hokkaido University.

### 2. Plasmids

Full-length YG1 S, M, and L segment constructs were generated by cloning into TVT7R (0,0), kindly provided by Dr. Benjamin Brennan, Glasgow Center for Virus Research, Scotland, United Kingdom (9). Cloning was conducted using an In-Fusion HD cloning kit (Takara Bio, CA, USA), and the resulting plasmids were named pTVT7_YG1_S, pTVT7_YG1_M, and pTVT7_YG1_L. The amino acid mutation Y328H was introduced into the pTVT7_YG1_M construct using site-directed mutagenesis using the KOD One PCR Master Mix (Toyobo, Osaka, Japan) and primers containing the mutation (Forward: CGTGTCAGACCAAAATGCCATGGTTTCTCCAGAATGA, and Reverse: TCATTCTGGAGAAACCATGGCATTTTGGTCTGACACG). The mutation N1891K was introduced to the pTVT7_YG1_L construct, and R624W was introduced to pTVT7_YG1_M using In-Fusion HD cloning kit (Takara Bio), primers containing mutation for insert sequence (5′-AACTTGGAAGTGCTTTGTGGTAGG-3′, 3′-GACCAGGCCCAATTGTCAAGAGTTTTC-3′ and primers containing mutation for vector sequence 5′-CAATTGGGCCTGGTCACATGCCTCAGTTC-3′, 5′-AAGCACTTCCAAGTTCATCTGGGCGTCT-3′).

### 3. Recombinant virus generation

Eight different recombinant viruses were generated by transfecting 4 × 10^5^ cells/mL BSR-T7/5 cells with 2 µg of each pTVT-based plasmids expressing viral genomic segments with or without point mutations (Table 1), 0.2 µg pTM1-HB29ppL and 1 µg pTM1-HB29N. Transfection experiments were conducted using TransIT-LT1 reagent (Mirus Bio LLC, Madison, WI, USA). After five days of incubation, the recombinant virus supernatant was harvested as P1. The P1 supernatant was then blindly passed into Vero E6 cells, and the recombinant virus stock P2 was prepared by harvesting the culture medium 7–9 days post-infection (dpi), depending on the CPE.

**Table 1.**
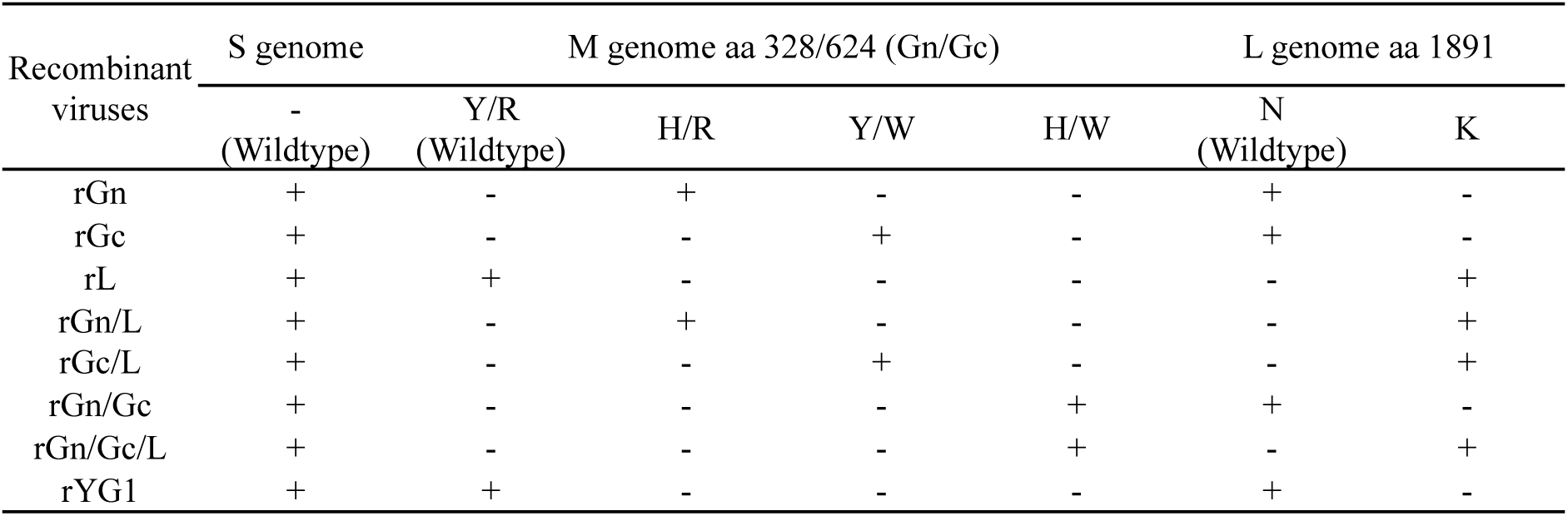
Recombinant viruses and the respective plasmid constructs used for the reverse genetics system.

The recombinant virus stock P2 was inoculated into Vero E6 cells for viral amplification, and the seed virus was obtained after six days of incubation. The eight rescued viruses were named after their mutations for ease of application (Table 1). Total RNA was extracted from the cells using Isogen (Nippon Gene, Tokyo, Japan), and the culture supernatant was extracted using Isogen LS (Nippon Gene) according to the manufacturer’s protocol. Real time PCR was performed using the ReverTra Ace qPCR RT Master Mix (Toyobo). The nucleotide sequences of the rescue viruses were determined using an amplicon-based ligation sequencing protocol with a native barcoding kit on a MinION device (SQK-LSK1-9 and EXP-NDB104; Oxford Nanopore Technologies, Oxford, UK) and Sanger sequencing to confirm that no mutations had occurred during rescue.

### 4. Virus titration using a plaque assay

Vero E6 cells were seeded at 4 × 10^5^ cells/mL in a 6-well plate, infected with serial dilutions of each recombinant virus, and cultured in an overlay medium: MEM containing 0.8% SeaKem ME agarose (Takara, Kusatsu, Japan) and 4% FBS. After seven dpi, 2 mL of neutral red solution (0.1 mg/mL in MEM; Sigma-Aldrich) was added to the overlay medium. After incubation for 24 h, viral foci were visible, and the number and diameter of the foci were examined. The titer of each recombinant virus was calculated.

### 5. Cell fusion assay

Vero E6 cells were seeded to a 96-well plate inoculated with serial dilutions of each recombinant virus and incubated for seven days at 37 °C. The medium was replaced with 50 mM acetate-buffered saline pH 5.6, and the cells were incubated for 2 min at room temperature. Then, the acetate buffer was replaced with a fresh growth medium and incubated for 24 h at 37 ^°^C. The cells were fixed with Mildform 10 NM, a 10% formalin neutral buffer-methanol solution (Wako Pure Chemical Corporation, Osaka, Japan) for 20 min and stained with azur-eosin-methylene blue solution (Muto Pure Chemicals Co., Ltd., Tokyo, Japan) to visualize cell fusion. Images were acquired using a KEYENCE/BZ-X800 microscope (Keyence Corporation, Osaka, Japan).

### 6. Pseudotype virus assay

The mammalian expression vector pCAGGS-YG1 GP and its mutants have been previously cloned (18). The pseudotype vesicular stomatitis virus (VSV), possessing green fluorescent protein and luciferase genes, was kindly provided by Dr. Heinz Feldmann (DIS, NIAID, NIH). Each pseudotyped VSV was generated as previously reported. Briefly, 293T cells were transfected with each plasmid vector, pCAG-GP-Y/R, pCAG-GP-H/R, pCAG-GP-Y/W, and pCAG-GP-H/W, using TransIT-LT1 (Mirus Bio LLC) and incubated at 37 °C for 30 h. Pseudotype VSV possessing VSV-G was inoculated, and the culture supernatant was collected 13 h after inoculation. The VSV particles produced in the collected supernatants were evaluated by western blotting using an anti-VSV M monoclonal antibody (kindly provided by Professor Ayato Takada, Hokkaido University). Western blotting was performed as described previously. Vero E6 cells were inoculated with ten times dilution of each VSVΔG/SFTSV (GP-Y/R, GP-H/R, GP-Y/W, or GP-H/W) and incubated at 37 °C for 18 h. Luciferase activity in the cell lysates was determined using the Dual-Glo luciferase assay system (Promega, Madison, WI, USA). The degree of cell entry was directly proportional to the *Renilla* luciferase units and normalized to firefly luciferase activity.

### 7. Treatments of a pan-caspase inhibitor

Vero E6 cells were inoculated with rGn/Gc/L, rL, and rYG1, and three days post-inoculation, media were replaced with fresh media containing 50 µM of pan-caspase inhibitor Z-VAD-FMK (Selleck chemicals, TX, USA). Dimethyl sulfoxide (Sigma-Aldrich) was used as a diluent for the pan-caspase inhibitors. Mock-infected Vero E6 cells were used as the controls. Five days after inoculation, the cells were fixed with MildForm (Wako Pure Chemical) and stained with Giemsa azure-eosin-methylene blue solution (Muto Pure Chemicals) for visualization.

### 8. Western blot analysis

To detect the programmed cell death factors in infected cells, the recombinant viruses, rL, rGn/Gc/L, and rYG1, were inoculated into Vero E6 cells in a 6-well plate at a multiplicity of infection 0.05. One- and four days post-inoculation, the cells were collected using a cell scraper and rinsed once with ice-cold phosphate-buffered saline (PBS), followed by lysis with 200 µL of sample buffer solution with 2-mercaptoethanol (Fujifilm-Wako) and incubated at 100 °C for 5 min. Furthermore, Vero E6 cells were stimulated with 500 ng/µL of actinomycin D (Sigma-Aldrich). After two days of stimulation, the cells were harvested, and the cell lysate was used as a positive control for apoptosis. U937 cells were stimulated with 250 nM phorbol-12-myristate-13-acetate for 16 h. Cells were primed with lipopolysaccharide (LPS) (200 ng/mL) for 4 h for differentiation into macrophage-like cells. Then, cells were stimulated with 7.5 μM nigericin for 6 h and used as the control for caspase induction using pyroptosis. Cells were lysed as above and incubated at 100 °C for 5 min. The lysates (15 µL) were loaded onto a sodium dodecyl-sulfate-polyacrylamide gel electrophoresis gel (e-PAGEL 1020 L, Atto, Tokyo, Japan) and electroblotted onto a 0.45-μm pore immunoblot polyvinylidene fluoride membrane (Millipore, Billerica, MA, USA). The membranes were incubated with primary antibodies: anti-caspase 1 (Abcam, Cambridge, UK) diluted 1,000 times, anti-poly (ADP-ribose) polymerase (PARP) (Cell Signaling, Danvers, MA, USA) diluted 1,000 times, and anti-caspase 3 (Sigma-Aldrich) diluted 1,00 times with Can Get Signal Immunoreaction Enhancer Solution 1 (Toyobo) for 1 h at room temperature.

Horseradish peroxidase-conjugated mouse anti-rabbit IgG (Jackson ImmunoResearch Laboratories Inc., Baltimore, MD, USA) diluted 10,000 times with Can Get Signal Immunoreaction Enhancer Solution 2 (Toyobo) was used as the secondary antibody and incubated for 1 h at room temperature. Bound antibodies reacted with Amersham ECL Prime (GE Healthcare Life Science, PA, USA) and were detected using ImageQuant LAS 4000 mini (GE Healthcare Life Science). An HRP-conjugated anti-glyceraldehyde-3-phosphate antibody (Proteintech, Rosemont, IL, USA) diluted 2,000 times with PBS was used as the housekeeping gene in this experiment.

### 9. Cell death inhibition in rYG1-infected cells

Actinomycin D (Sigma-Aldrich) in high to low concentrations (500, 400, 300, 200, and 100 ng/µL) were mixed with equal volumes of rYG1 with 10 plaque-forming units (PFU)/well and inoculated into the monolayer of Vero E6, Huh-7, and BHK/T7-9 cells. rYG1 infection and actinomycin D stimulation alone were used as controls. After three days of incubation, the cells were fixed and stained as described above.

### 10. Co-infection of recombinant viruses

The recombinant wild-type virus rYG1 was inoculated into Vero E6 cells at a multiplicity of infection of 0.2 together with rGn, rGc, and rL viruses (ratios of 10:1. 1:1, and 0.1:1) were inoculated into Vero E6 cells at a gradient dilution in a monolayer 6-well plate. Eight days after the inoculation, the culture supernatant was collected and centrifuged to remove the cells. Total RNA was extracted from the culture supernatant using Isogen LS (Nippon Gene), according to the manufacturer’s instructions, and cDNA was produced using a ReveTra Ace cDNA synthesis kit (Nippon Genetics Co., Ltd, Tokyo, Japan). To determine the dominant virus in each fraction, M genome segment 804–1325, 1792–2381, and L genome segment 5135–5883 were amplified and sequenced using MinION, as described above, and the ratio of wild-type virus to mutant virus was estimated.

### 11. Statistical analysis

Student’s t-test was conducted in plaque size and luciferase activity comparisons.

## Results

### 1. Establishment of recombinant viruses using a reverse genetics system

We established a reverse genetics system for the SFTSV-YG1 strain and rescued seven recombinant viruses of the original YG1 strain with single, double, or triple mutations using the same method. Together with helper plasmids, as shown in Table 1, the transcription plasmids were transfected into BSR-T7/5 cells in a 60-mm culture dish. The supernatant was harvested after five days of incubation, and the P1 virus was obtained. The P1 virus was inoculated into Vero E6 cells for expansion, and P2 was obtained as the seed virus at 7–9 dpi. The rescued viruses were designated as rGn, rGc, rL, rGn/L, rGc/L, rGn/Gc, rGn/Gc/L, or rYG1, depending on the mutation(s) (Table 1). We detected the genomes of the respective recombinant viruses using the P2 viruses used for genome detection via real time PCR and indirect immunofluorescence antibody tests (data not shown). Further passaging was conducted using Vero E6 cells to obtain the P3 virus, which was used for downstream analysis as the working virus. The nucleotide sequences of all recombinant viruses were determined and confirmed using MinION and Sanger sequencing, and there were no unnecessary mutations (data not shown).

### 2. Plaque formation and titers of recombinant viruses

As shown in Fig. 1, all recombinant viruses other than rYG1 showed clear plaques, and virus titers were higher than 10^6^ PFU/mL, except for rGn/Gc/L. The plaques formed by the recombinant viruses revealed different phenotypes, depending on the mutations in the virus. Viral plaques caused by rGn have a clear, smooth, and large appearance, whereas those caused by rGc have a large but rough and smoky appearance. In contrast, rGn/L produced the largest number of plaques. Gn and Gc mutations have a synergistic effect on increasing plaque size. In contrast, rL plaques appeared as pinpricks. The cells in the rL plaque had died. The plaques of rYG1 were similar in size to those of rL but were very smoky, which was difficult to see in the picture. With the addition of the rL mutation, the cells in the plaque were killed so that the plaque appeared transparent.

**Figure 1:**
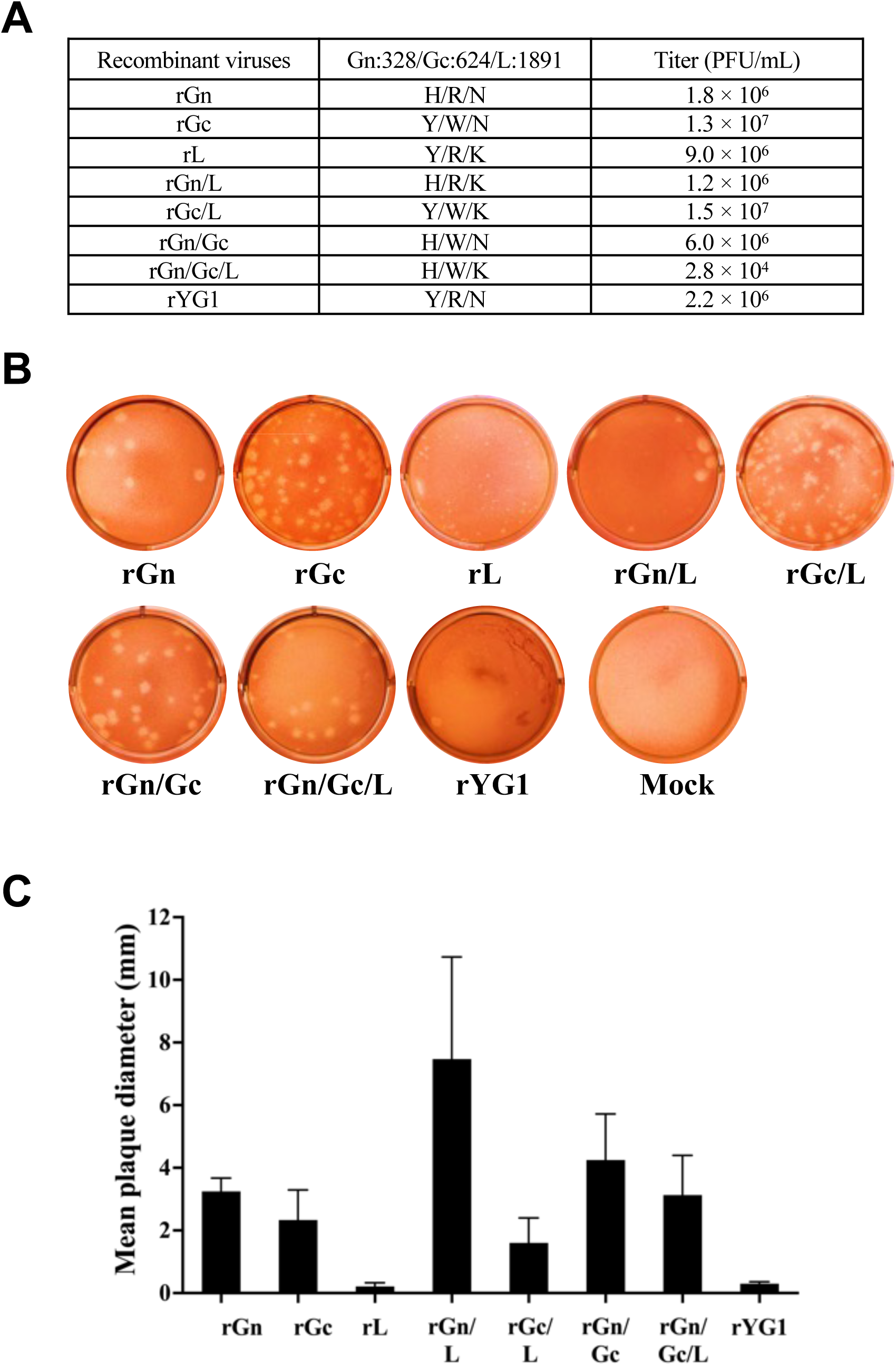
Virus titration and plaque-forming activity of recombinant viruses. A; Recombinant viruses and their titers of working stock. B; Plaques of the recombinant viruses. Plaque photographs were taken 10 days after inoculation and three days after the overlay of neutral red, except for rGc and rGn/Gc, which were taken nine days after infection. C; Size of plaques. The plaque diameters were measured. The results show the mean and standard deviation of 5-25 plaques.

### 3. Assessment of low pH-dependent cell fusion activity and CPE of recombinant viruses

In a previous study, we demonstrated that a mutation in R624W in Gc induced strong syncytia formation under low pH conditions, and subclone B7 only demonstrated robust CPE in infected cells. The recombinant virus rGc showed strong syncytial formation under low pH conditions. rGn showed weak cell fusion activity compared with rGc under low-pH conditions. In contrast, rL showed intense CPE under low and normal pH conditions but no cell fusion activity (Fig. 2). Both double mutant viruses, rGn/L and rGc/L, exhibited intense CPE and cell fusion activity. However, the robust CPE made it challenging to observe low pH-dependent cell fusion activity, owing to the detachment of cells (Fig. 2). These results suggested that N1891K is essential for inducing CPE, whereas R624W is involved in cell fusion.

**Figure 2:**
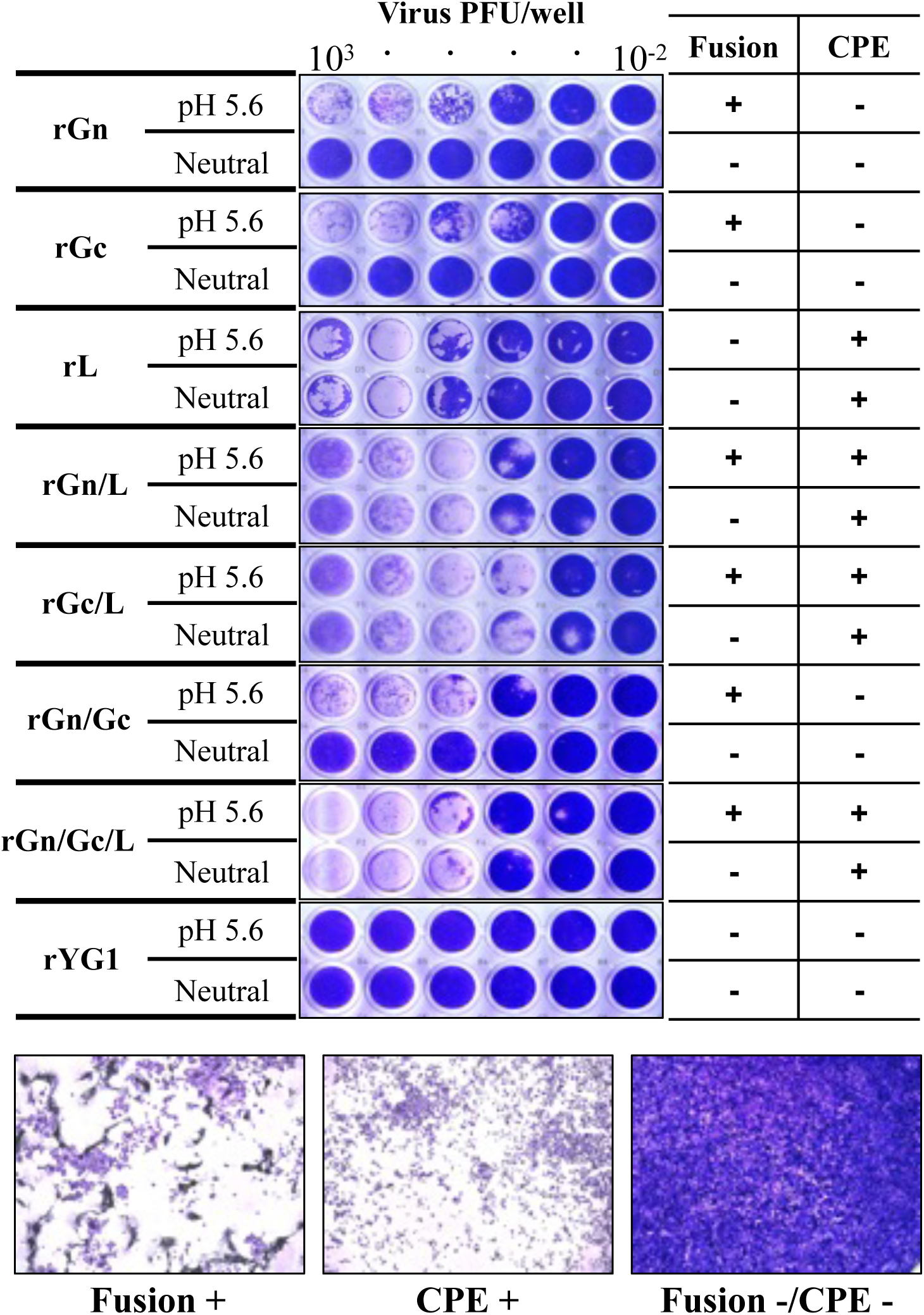
Assessment of low pH-dependent cell fusion activity and CPE of recombinant viruses Vero E6 cells were inoculated with serial dilution of each recombinant virus and incubated for seven days at 37 °C. The medium was replaced with 50 mM acetate-buffered saline (pH 5.6), and the cells were incubated for 2 min at room temperature. Then, the acetate buffer was replaced with fresh growth medium and incubated for 24 h at 37 °C, 5% CO_2_. The wells in the upper panels show the Giemsa staining results after low-pH treatment. In clear wells, numerous syncytia were found by cell fusion. Wells in the lower panels show Giemsa staining results without low pH treatment. The cells were washed with CPE. Cells were stained without cell fusion or CPE. Examples of the evaluations are shown below.

### 4. Effects of amino acid alteration on GP on the step of virus entry to Vero cells

To examine the efficacy of viral cell entry, we generated pseudotyped viruses bearing YG1-GP-Y/R, GP-H/R, GP-Y/W, or GP-H/W, inoculated them into Vero E6 cells, and measured luciferase activity. The luciferase activity of VSVΔG/SFTSV-GP-H/R was significantly lower than that of VSVΔG/SFTSV -GP-Y/R (Fig. 3). Luciferase activity of VSVΔG/SFTSV-GP-H/W having both mutations (Y328H and R624W) was recovered to that of wild-type GP. The mutation in Gc alone, VSVΔG/SFTSV - GP-Y/W, did not show any significant difference, meaning that the mutation R624W does not have a particular effect on viral entry alone.

**Figure 3:**
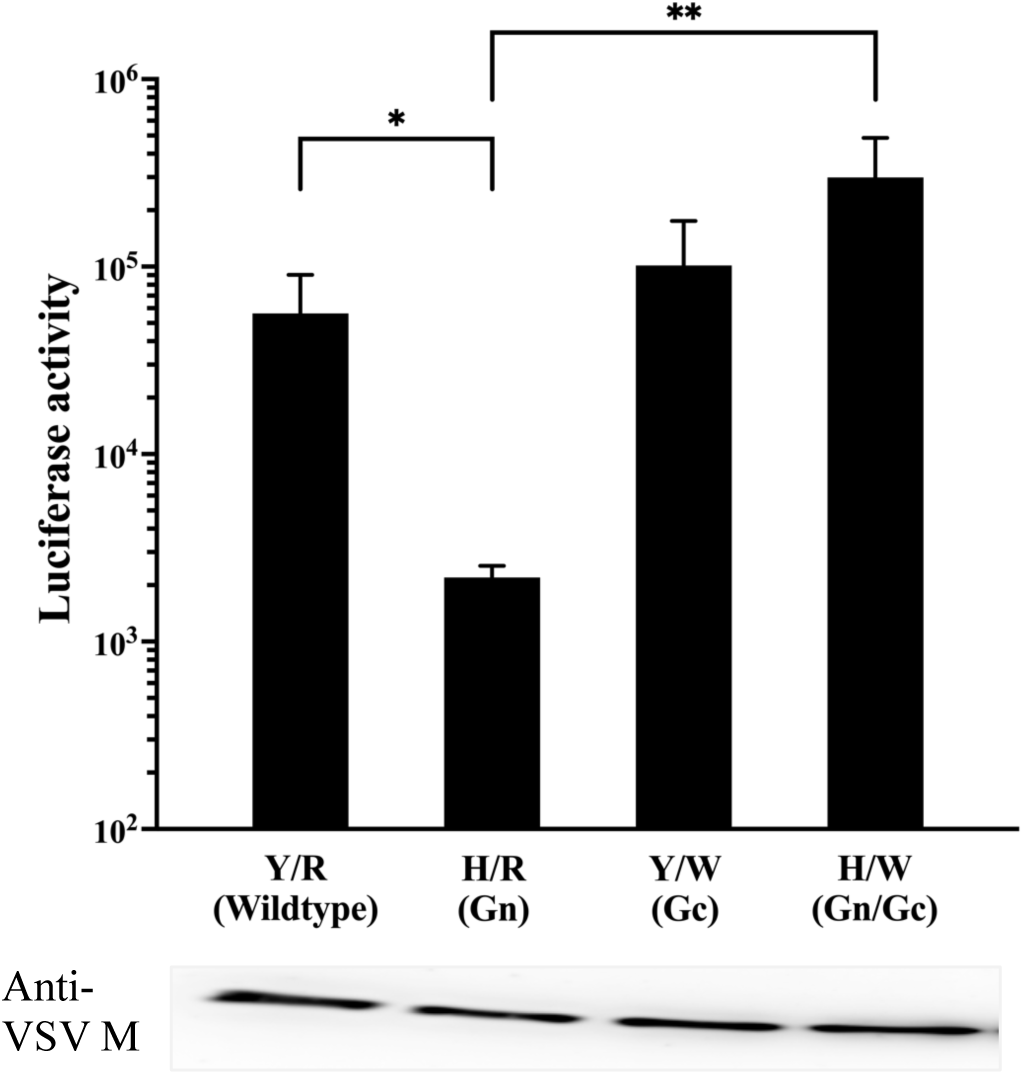
Comparison of viral entry using pseudotype viruses bearing wild-type GP of YG1 or GPs with single/double mutations at 328 and 624. Pseudotyped vesicular stomatitis viruses bearing GP-Y/R, GP-H/R, GP-Y/W, or GP-H/W instead of VSV-G were inoculated to the Vero E6 cells, and the luciferase activity was measured at 18 h post-inoculation. The approximate viral inoculum was evaluated using western blotting using an anti-VSV-M antibody. Results are presented as means and standard deviations of triplicate luciferase activity experiments.

### 5. Programmed cell death related to SFTSV infection

We examined the effects of the pan-caspase inhibitor Z-VAD-FMK on rGn/Gc/L-, rL-, and rYG1-infected cells. As shown in Fig. 4, Z-VAD-FMK reduced cell death induced by rGn/Gc/L and rL infection compared with that in cells without inhibitors. In contrast, rYG1 infection did not cause cell death with or without an inhibitor, indicating that the cell death induced by rGn/Gc/L and rL was caspase-dependent. Next, we evaluated the expression of programmed cell death-related molecules in SFTSV-infected cells. PARP cleavage was observed in the apoptosis-induced Vero E6 cells treated with actinomycin D (Fig. 5A). In contrast, cleaved PARP bands were not detected in the rL-, rGn/Gc/L-, or mock-infected cells at any time point (1 or 4 dpi). Programmed cell death induced by rL and rGn/Gc/L infection might not be due to apoptosis but due to another mechanism. Then, we examined the induction of caspases 1 and 3. As shown in Fig. 5A and 5C, caspases 1 and 3 were detected in pyroptosis-induced U937 cells treated with nigericin. Caspase induction at 4 dpi was observed in both CPE-inducing (rL and rGn/Gc/L) and CPE-non-inducing (rYG1) virus-inoculated cells. In contrast, Caspase 1 was not detected following actinomycin D stimulation (Figure 5B). These results suggest that both CPE-inducing and non-inducing SFTSV trigger programmed cell death, which may be pyroptosis.

**Figure 4:**
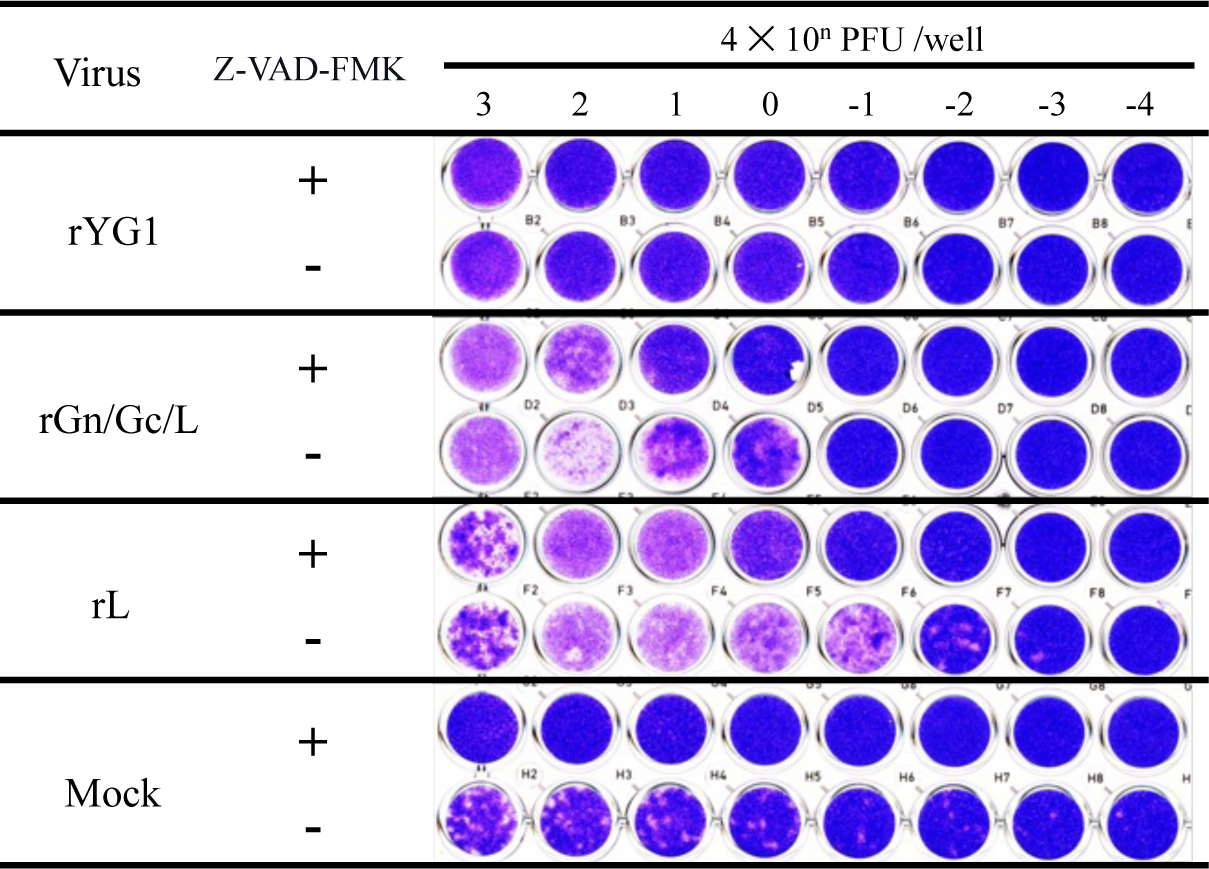
Inhibitory effects of pan-caspase inhibitor on CPE induced by recombinant viruses. Vero E6 cells were inoculated with recombinant viruses, and media were replaced with fresh media, 50 µM caspase inhibitor, Z-VAD-FMK in dimethyl sulfoxide (+) or dimethyl sulfoxide alone as diluent control (-) at three days post-inoculation. Mock-infected Vero E6 cells were used as the controls. Five days post-inoculation, the cells were fixed and stained with Giemsa azure-eosin-methylene blue solution.

**Figure 5:**
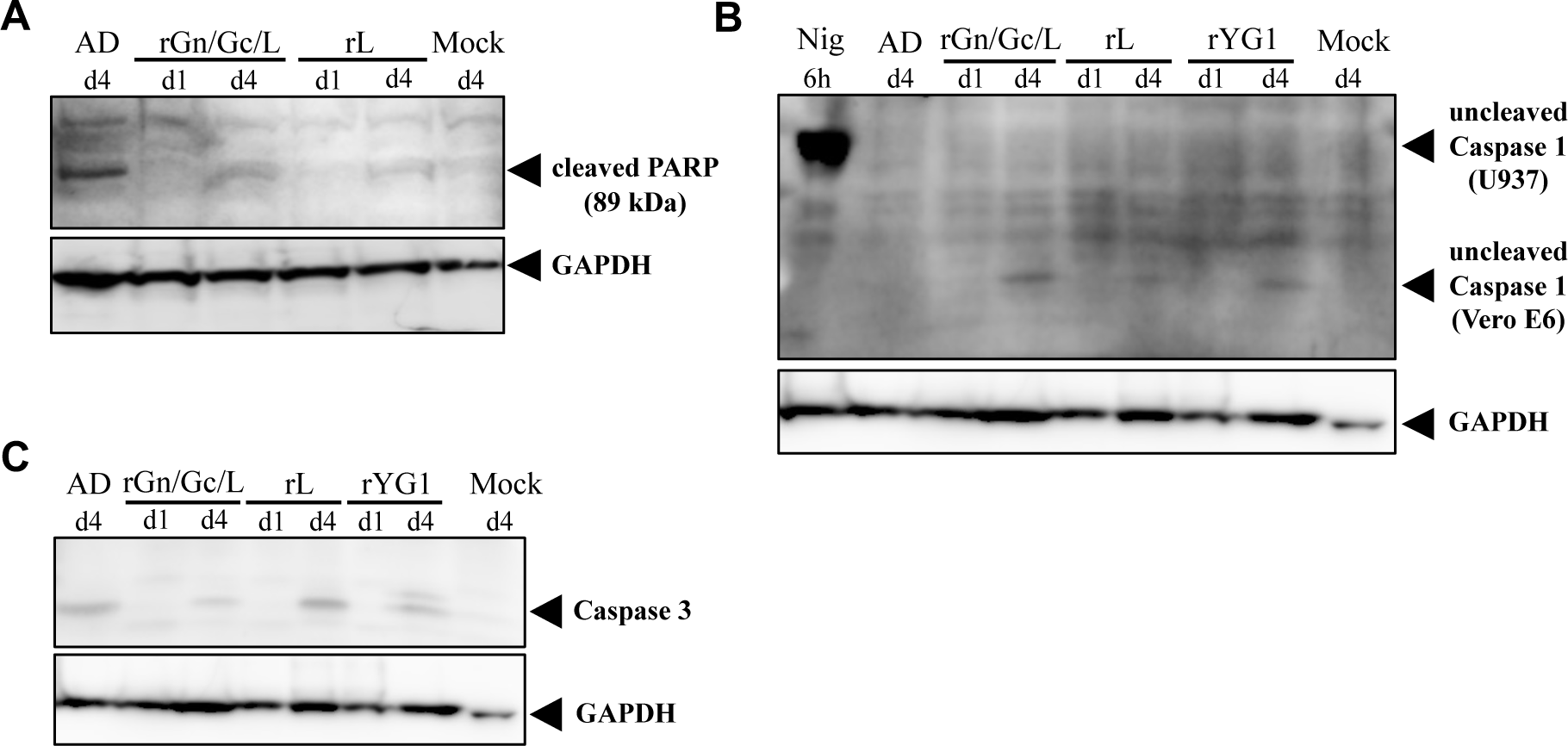
Induction of cleaved PARP, caspase-1, and caspase-3 in SFTSV-infected cells using western blot assay Recombinant viruses rL, rGn/Gc/L, and rYG1 were inoculated into Vero E6 cells. One and four days after inoculation, the cells were harvested and examined. As the positive control of programmed cell death, Vero E6 cells were stimulated with 500 ng/µL of actinomycin D. U937 cells were stimulated by phorbol-12-myristate-13-acetate for 16 h and primed with LPS for 4 h to differentiate them into macrophage-like cells. Then, differentiated U937 cells were stimulated with 7.5 μM nigericin for 6 h and used as the control for caspase-1 induction by pyroptosis. Glyceraldehyde-3-phosphate was used as the standard control. A: The induction of cleaved PARP was used as an apoptotic marker. The actinomycin D-treated Vero E6 cell lysate was used as a positive control for cleaved PARP. B: Caspase-1 induction was used as a pyroptosis marker. Nigericin-treated U937 cell lysates were used as a control for caspase-1 induction. C: Caspase-3 is an important mediator of programmed cell death. Actinomycin D-treated Vero E6 cell lysate was used as a positive control for caspase-3 induction.

### 6. Inhibitory effect of wild-type SFTSV on actinomycin D-induced cell death

Actinomycin D, an apoptosis inducer, induces programmed cell death in Vero E6, BHK-T7/9, and Huh-7 cells. As shown in Fig. 6, rYG1 infection alone did not induce any cell damage. Furthermore, rYG1 inoculation reduced actinomycin D-induced cell damage. Although actinomycin D-treated cells exhibited dose-dependent damage, high concentrations of actinomycin D-induced cell death in rYG1- and mock-infected cells. As the actinomycin D concentration decreased, cell death was suppressed by rYG1 in all cell lines. These results suggested that rYG1 infection inhibited programmed cell death induced by actinomycin D.

**Figure 6:**
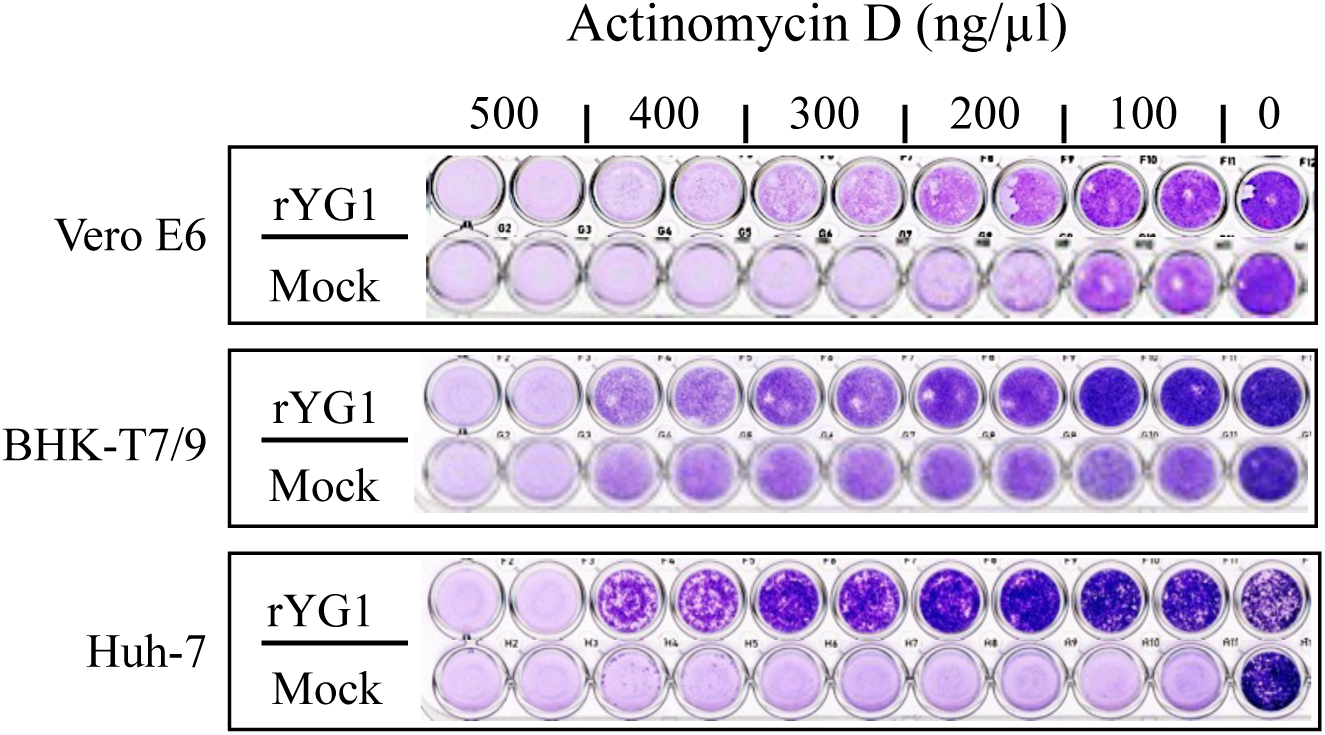
The impact of SFTSV on actinomycin D-induced cell death in line cells. Vero E6, Huh-7, and BHK/T7-9 cell monolayers were inoculated with 10 PFU of the rYG1 virus per well. Actinomycin D in high to low concentrations (500, 400, 300, 200, and 100 ng/µL) were mixed with inoculum before inoculation. rYG1 infection and actinomycin D stimulation alone (mock infection) were used as controls. After three days of incubation, the cells were fixed and stained with Giemsa azure-eosin-methylene blue solution for visualization.

### 7. Competition of wild-type and recombinant viruses in virus propagation

Next, we evaluated the co-infection with rYG1 and recombinant viruses. Monolayered Vero E6 cells in a 6-well plate were co-inoculated with rYG1 and rGn, rGc, or rL. After 1 h of incubation, each supernatant was replaced with the growth medium and incubated for seven days. The ratio of rYG1 to recombinant viruses in the culture supernatant was estimated (Table 2). Even in 10 times higher inoculation dose, recombinant viruses were not the major population in the culture supernatant. In the 1:10 rL inoculation, the mutant virus population was reduced to 21.7% after seven days of infection. In addition, CPE were not observed for any combination.

**Table 2.**
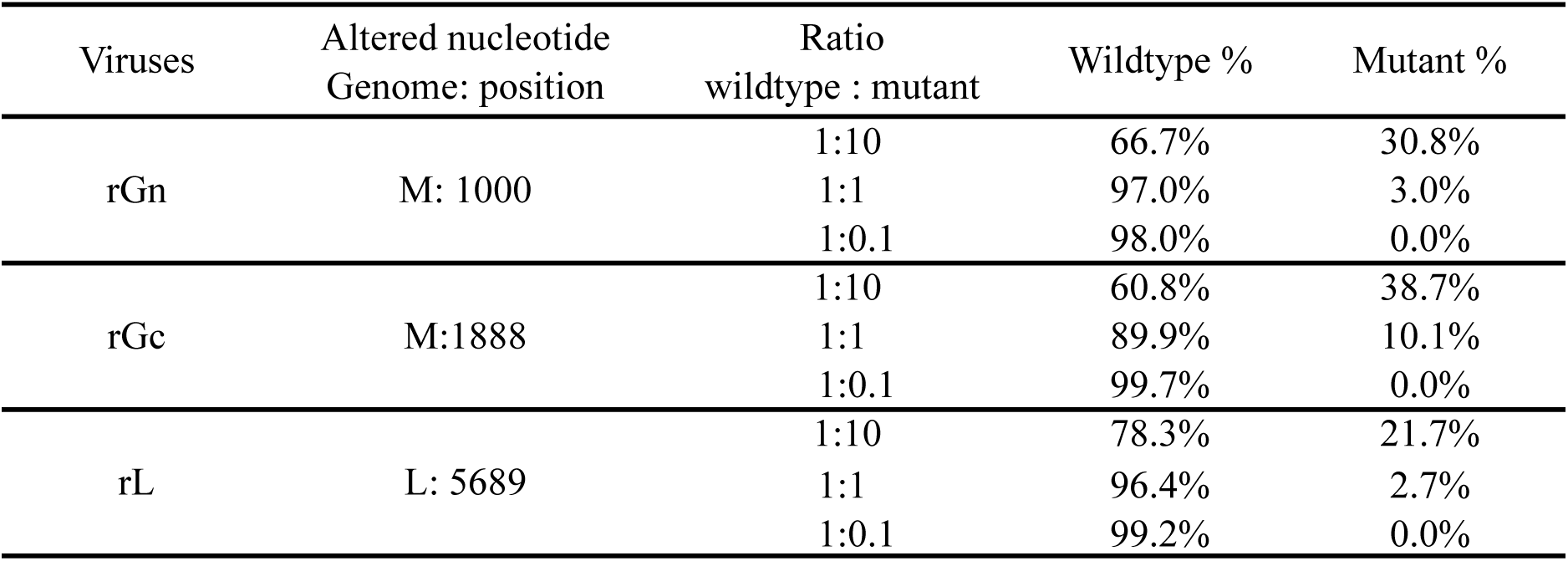
Viral propagation in co-infection with recombinant viruses.

## Discussion

SFTS is an emerging infectious disease with a high fatality rate caused by a member of the *Pheniuviridae* family, the *Bandavirus* genus, which has become a significant health concern in most Asian countries owing to the lack of therapeutic and pathological knowledge (26–30). Several studies have reported the importance of quasispecies in the pathogenicity and growth strategies of various viruses (31, 32). Understanding the characteristics of the constructs in SFTSV quasispecies may shed light on viral propagation and the pathogenicity of the population.

Our previous findings showed that the amino acid at position 624 in Gc of the original YG1 strain of SFTSV plays an essential role in low pH-dependent cell fusion activity (18). In addition, the polarity of amino acid position 1891 in the L protein of YG1 is critical for polymerase activity, and the L-segment C-terminal domain is critical for genome transcription and viral replication (23). However, the role of the Y328H variation in Gn is yet to be clarified. The Y328H variation in Gn was found in nearly 30% of the viral population in patient blood, but after virus isolation by Vero E6 cells, the ratio of this Gn variant was reduced (24). This explains why SFTSV in the patient’s blood consisted of almost 70% rYG1 and 30% rGn, which are highly pathogenic species in this fatal infection. It is assumed that a selective pressure exists in patients with the Y328H mutation in Gn to become a quasispecies. To detect the virological characteristics of each mutation in the YG1 quasispecies *in vitro*, we recovered recombinant viruses using a newly developed reverse genetics system for the YG1 strain. Data obtained from the plaque-forming assay revealed phenotypic differences in the foci of each recombinant virus. In this study, the rGn virus with the Y328H alteration in Gn showed large-sized plaques, suggesting that there may be differences in the virus spreading patterns from infected cells to neighboring cells in an overlay medium. Furthermore, the pseudotype assay in the current study showed that the Y328H mutation in GP discouraged cell entry, and the mutation in Gc (R624W) significantly compensated for this disadvantage in the entry step. This can be explained by the simultaneous increase in the growth of Vero E6 cells, which is more advantageous in the entry step. This observation suggests that the Gn variant may have acquired R624W Gc mutations during adapting to Vero E6 cells.

SFTSV Gn binds to non-muscle myosin heavy chain IIA, which is critical for the cellular entry of SFTSV (33). A recent study has shown that phlebovirus Gc plays a vital role in the fusion process. In contrast, Gn likely contributes to receptor binding depending on its position on the viral surface (34). The Y328H variation was disadvantageous for the entry to Vero E6 but might be advantageous for unknown target cells *in vivo*. The Gc structure of Rift Valley fever virus is similar to that of the class II fusion protein architecture, which has been identified as an effector of membrane fusion (35). In this study, the recombinant viruses with R624W alterations showed high fusion activity. However, we did not find subclones with only R624W alterations in Gc throughout the limiting dilution cloning process. Altogether, with this observation and the results of the competitive growth assay, the virus with the R624W alteration alone was also disadvantageous for the growth of Vero E6 cells.

Alteration of N1891K in the L protein plays an essential role in CPE induction. However, the mechanisms underlying the induction of cell death remain unclear. A recent study suggested that during the SFTSV infection, it activates the nucleotide-binding domain, leucine-rich– containing family, pyrin domain–containing-3 inflammasome and pyroptosis, leading to IL-1β/IL-18 secretion (21, 36–39). The necrotic cell death inhibitor, IM-54, did not affect cell death caused by B7 or rL (data not shown). The pan-caspase inhibitor Z-VAD-FMK inhibited cell death induced by rL and rGn/Gc/L, indicating a caspase-dependent mechanism (Fig. 4). Herein, we can rule out the possibility of apoptosis since cleavage of the PARP antibody by actinomycin D treatment was not prominent in rL compared with apoptosis-infected cells. Both CPE-inducing and non-inducing viruses induced caspase 1 and 3. Wild-type viruses allow cells to escape death after the induction of caspases 1 and 3. Only viruses with the N1891K mutation in the L genome segment failed to suppress cell death for unknown reasons. Suppression of cell death by SFTSV was observed during apoptotic cell death induced by actinomycin D. SFTSV probably acts close to the final stage of cell death, which is common in pyroptosis and apoptosis.

We previously reported that the alteration of N1891K in the L protein induced high RdRp activity (23). However, the relationship between the high RdRp activity and cell death remains unclear. One possibility is that the high production of viral RNA and proteins within the cell causes significant stress, leading to cell death. Another possibility is that the L protein directly suppresses cell death, and the N1891K portion is involved in this step. Further studies are required to clarify the mechanism underlying CPE caused by this alteration. However, this variation was not advantageous for viral survival in Vero cells. The wild-type virus easily outperformed the rL virus. Since the induction of CPE in infected cells is not thought to be beneficial for viral survival, SFTSV might have a mechanism to avoid cell death. When a cell is infected with a virus, whether it produces interferon to be antiviral or whether it chooses cell death and eliminates itself is based on the same mechanism mediated by apoptosis signal-regulating kinase family kinases and is selected depending on the situation (40). During SFTSV infection, nonstructural S proteins play a role in suppressing the interferon system (41–45). This study shows that SFTSV may also have a mechanism to resist cell death.

In this study, we obtained information on viral pathogenesis by analyzing quasispecies derived from one fatal case. This suggests that combining mutations may increase the viability of mutant viruses, selecting viruses to create a suitable population for propagation, leading to the appearance of new pathogenic viruses.

## References

1. Xu B, Liu L, Huang X, Ma H, Zhang Y, Du Y, Wang P, Tang X, Wang H, Kang K, Zhang S, Zhao G, Wu W, Yang Y, Chen H, Mu F, Chen W. 2011. Metagenomic analysis of fever, thrombocytopenia and leukopenia syndrome (FTLS) in Henan Province, China: discovery of a new bunyavirus. PLoS Pathog 7:e1002369.

2. Takahashi T, Maeda K, Suzuki T, Ishido A, Shigeoka T, Tominaga T, Kamei T, Honda M, Ninomiya D, Sakai T, Senba T, Kaneyuki S, Sakaguchi S, Satoh A, Hosokawa T, Kawabe Y, Kurihara S, Izumikawa K, Kohno S, Azuma T, Suemori K, Yasukawa M, Mizutani T, Omatsu T, Katayama Y, Miyahara M, Ijuin M, Doi K, Okuda M, Umeki K, Saito T, Fukushima K, Nakajima K, Yoshikawa T, Tani H, Fukushi S, Fukuma A, Ogata M, Shimojima M, Nakajima N, Nagata N, Katano H, Fukumoto H, Sato Y, Hasegawa H, Yamagishi T, Oishi K, Kurane I, Morikawa S, Saijo M. 2014. The first identification and retrospective study of Severe Fever with Thrombocytopenia Syndrome in Japan. J Infect Dis 209:816–27.

3. Kim KH, Ko MK, Kim N, Kim HH, Yi J. 2017. Seroprevalence of Severe Fever with Thrombocytopenia Syndrome in Southeastern Korea, 2015. J Korean Med Sci 32:29–32.

4. Tran XC, Yun Y, Van An L, Kim SH, Thao NTP, Man PKC, Yoo JR, Heo ST, Cho NH, Lee KH. 2019. Endemic Severe Fever with Thrombocytopenia Syndrome, Vietnam. Emerg Infect Dis 25:1029–1031.

5. Liu Q, He B, Huang SY, Wei F, Zhu XQ. 2014. Severe fever with thrombocytopenia syndrome, an emerging tick-borne zoonosis. Lancet Infect Dis 14:763–72.

6. Robles NJC, Han HJ, Park SJ, Choi YK. 2018. Epidemiology of severe fever and thrombocytopenia syndrome virus infection and the need for therapeutics for the prevention. Clin Exp Vaccine Res 7:43–50.

7. Kobayashi Y, Kato H, Yamagishi T, Shimada T, Matsui T, Yoshikawa T, Kurosu T, Shimojima M, Morikawa S, Hasegawa H, Saijo M, Oishi K, Japan SERG. 2020. Severe Fever with Thrombocytopenia Syndrome, Japan, 2013-2017. Emerg Infect Dis 26:692–699.

8. Mita T. 2019. EpidemiologyofSevereFeverwithThrombocytopeniaSyndromeinJapan. Juntendo Medical Journal 65:130–135.

9. Brennan B, Li P, Zhang S, Li A, Liang M, Li D, Elliott RM. 2015. Reverse genetics system for severe fever with thrombocytopenia syndrome virus. J Virol 89:3026–37.

10. Elliott RM, Schmaljohn CS. 2013. Bunyaviridae, p 1244–1282. In Knipe DM, Howley PM (ed), Fields Virology, 6th ed. Wolters Kluwerz.

11. Tani H. 2014. Analyses of Entry Mechanisms of Novel Emerging Viruses Using Pseudotype VSV System. Trop Med Health 42:71–82.

12. Plegge T, Hofmann-Winkler H, Spiegel M, Pohlmann S. 2016. Evidence that Processing of the Severe Fever with Thrombocytopenia Syndrome Virus Gn/Gc Polyprotein Is Critical for Viral Infectivity and Requires an Internal Gc Signal Peptide. PLoS One 11:e0166013.

13. Hofmann H, Li X, Zhang X, Liu W, Kuhl A, Kaup F, Soldan SS, Gonzalez-Scarano F, Weber F, He Y, Pohlmann S. 2013. Severe fever with thrombocytopenia virus glycoproteins are targeted by neutralizing antibodies and can use DC-SIGN as a receptor for pH-dependent entry into human and animal cell lines. J Virol 87:4384–94.

14. Tani H, Shimojima M, Fukushi S, Yoshikawa T, Fukuma A, Taniguchi S, Morikawa S, Saijo M. 2016. Characterization of Glycoprotein-Mediated Entry of Severe Fever with Thrombocytopenia Syndrome Virus. J Virol 90:5292–5301.

15. Filone CM, Heise M, Doms RW, Bertolotti-Ciarlet A. 2006. Development and characterization of a Rift Valley fever virus cell-cell fusion assay using alphavirus replicon vectors. Virology 356:155–64.

16. Lozach PY, Mancini R, Bitto D, Meier R, Oestereich L, Overby AK, Pettersson RF, Helenius A. 2010. Entry of bunyaviruses into mammalian cells. Cell Host Microbe 7:488–99.

17. Tani H, Kawachi K, Kimura M, Taniguchi S, Shimojima M, Fukushi S, Igarashi M, Morikawa S, Saijo M. 2019. Identification of the amino acid residue important for fusion of severe fever with thrombocytopenia syndrome virus glycoprotein. Virology 535:102–110.

18. Tsuda Y, Igarashi M, Ito R, Nishio S, Shimizu K, Yoshimatsu K, Arikawa J. 2017. The amino acid at position 624 in the glycoprotein of SFTSV (severe fever with thrombocytopenia virus) plays a critical role in low-pH-dependent cell fusion activity. Biomed Res 38:89–97.

19. Nishio S, Tsuda Y, Ito R, Shimizu K, Yoshimatsu K, Arikawa J. 2017. Establishment of Subclones of the Severe Fever with Thrombocytopenia Syndrome Virus YG1 Strain Selected Using Low pH-Dependent Cell Fusion Activity. Jpn J Infect Dis 70:388–393.

20. Gowen BB, Westover JB, Miao J, Van Wettere AJ, Rigas JD, Hickerson BT, Jung KH, Li R, Conrad BL, Nielson S, Furuta Y, Wang Z. 2017. Modeling Severe Fever with Thrombocytopenia Syndrome Virus Infection in Golden Syrian Hamsters: Importance of STAT2 in Preventing Disease and Effective Treatment with Favipiravir. J Virol 91.

21. Gao C, Yu Y, Wen C, Li Z, Ding H, Qi X, Cardona CJ, Xing Z. 2022. Nonstructural Protein NSs Activates Inflammasome and Pyroptosis through Interaction with NLRP3 in Human Microglial Cells Infected with Severe Fever with Thrombocytopenia Syndrome Bandavirus. J Virol 96:e0016722.

22. Suzuki T, Sato Y, Sano K, Arashiro T, Katano H, Nakajima N, Shimojima M, Kataoka M, Takahashi K, Wada Y, Morikawa S, Fukushi S, Yoshikawa T, Saijo M, Hasegawa H. 2020. Severe fever with thrombocytopenia syndrome virus targets B cells in lethal human infections. J Clin Invest 130:799–812.

23. Noda K, Tsuda Y, Kozawa F, Igarashi M, Shimizu K, Arikawa J, Yoshimatsu K. 2020. The Polarity of an Amino Acid at Position 1891 of Severe Fever with Thrombocytopenia Syndrome Virus L Protein Is Critical for the Polymerase Activity. Viruses 13.

24. Yoshikawa T, Shimojima M, Fukushi S, Tani H, Fukuma A, Taniguchi S, Singh H, Suda Y, Shirabe K, Toda S, Shimazu Y, Nomachi T, Gokuden M, Morimitsu T, Ando K, Yoshikawa A, Kan M, Uramoto M, Osako H, Kida K, Takimoto H, Kitamoto H, Terasoma F, Honda A, Maeda K, Takahashi T, Yamagishi T, Oishi K, Morikawa S, Saijo M. 2015. Phylogenetic and Geographic Relationships of Severe Fever With Thrombocytopenia Syndrome Virus in China, South Korea, and Japan. J Infect Dis 212:889–98.

25. Buchholz UJ, Finke S, Conzelmann KK. 1999. Generation of bovine respiratory syncytial virus (BRSV) from cDNA: BRSV NS2 is not essential for virus replication in tissue culture, and the human RSV leader region acts as a functional BRSV genome promoter. J Virol 73:251–9.

26. Miyauchi A, Sada KE, Yamamoto H, Iriyoshi H, Touyama Y, Hashimoto D, Nojima S, Yamanaka S, Ishijima K, Maeda K, Kawamura M. 2022. Suspected Transmission of Severe Fever with Thrombocytopenia Syndrome Virus from a Cat to a Veterinarian by a Single Contact: A Case Report. Viruses 14.

27. Yu XJ, Liang MF, Zhang SY, Liu Y, Li JD, Sun YL, Zhang L, Zhang QF, Popov VL, Li C, Qu J, Li Q, Zhang YP, Hai R, Wu W, Wang Q, Zhan FX, Wang XJ, Kan B, Wang SW, Wan KL, Jing HQ, Lu JX, Yin WW, Zhou H, Guan XH, Liu JF, Bi ZQ, Liu GH, Ren J, Wang H, Zhao Z, Song JD, He JR, Wan T, Zhang JS, Fu XP, Sun LN, Dong XP, Feng ZJ, Yang WZ, Hong T, Zhang Y, Walker DH, Wang Y, Li DX. 2011. Fever with thrombocytopenia associated with a novel bunyavirus in China. N Engl J Med 364:1523–32.

28. Gong Z, Gu S, Zhang Y, Sun J, Wu X, Ling F, Shi W, Zhang P, Li D, Mao H, Zhang L, Wen D, Zhou B, Zhang H, Huang Y, Zhang R, Jiang J, Lin J, Xia S, Chen E, Chen Z. 2015. Probable aerosol transmission of severe fever with thrombocytopenia syndrome virus in southeastern China. Clin Microbiol Infect 21:1115–20.

29. Choi SJ, Park SW, Bae IG, Kim SH, Ryu SY, Kim HA, Jang HC, Hur J, Jun JB, Jung Y, Chang HH, Kim YK, Yi J, Kim KH, Hwang JH, Kim YS, Jeong HW, Song KH, Park WB, Kim ES, Oh MD, for Korea SCN. 2016. Severe Fever with Thrombocytopenia Syndrome in South Korea, 2013-2015. PLoS Negl Trop Dis 10:e0005264.

30. Kato H, Yamagishi T, Shimada T, Matsui T, Shimojima M, Saijo M, Oishi K, group-Japan Ser. 2016. Epidemiological and Clinical Features of Severe Fever with Thrombocytopenia Syndrome in Japan, 2013-2014. PLoS One 11:e0165207.

31. Grande-Pérez A, Martin V, Moreno H, de la Torre JC. 2015. Quasispecies: From Theory to Experimental Systems. Current Topics in Microbiology and Immunology. *In* Domingo E, Schuster P (ed), vol 392. Springer Cham.

32. Sevilla N, de la Torre JC. 2006. Arenavirus diversity and evolution: quasispecies in vivo. Curr Top Microbiol Immunol 299:315–35.

33. Sun Y, Qi Y, Liu C, Gao W, Chen P, Fu L, Peng B, Wang H, Jing Z, Zhong G, Li W. 2014. Nonmuscle myosin heavy chain IIA is a critical factor contributing to the efficiency of early infection of severe fever with thrombocytopenia syndrome virus. J Virol 88:237–48.

34. Wu Y, Zhu Y, Gao F, Jiao Y, Oladejo BO, Chai Y, Bi Y, Lu S, Dong M, Zhang C, Huang G, Wong G, Li N, Zhang Y, Li Y, Feng WH, Shi Y, Liang M, Zhang R, Qi J, Gao GF. 2017. Structures of phlebovirus glycoprotein Gn and identification of a neutralizing antibody epitope. Proc Natl Acad Sci U S A 114:E7564–E7573.

35. Dessau M, Modis Y. 2013. Crystal structure of glycoprotein C from Rift Valley fever virus. Proc Natl Acad Sci U S A 110:1696–701.

36. Li Z, Hu J, Bao C, Gao C, Zhang N, Cardona CJ, Xing Z. 2022. Activation of the NLRP3 inflammasome and elevation of interleukin-1beta secretion in infection by sever fever with thrombocytopenia syndrome virus. Sci Rep 12:2573.

37. Liu JW, Chu M, Jiao YJ, Zhou CM, Qi R, Yu XJ. 2021. SFTSV Infection Induced Interleukin-1beta Secretion Through NLRP3 Inflammasome Activation. Front Immunol 12:595140.

38. Li S, Li H, Zhang YL, Xin QL, Guan ZQ, Chen X, Zhang XA, Li XK, Xiao GF, Lozach PY, Cui J, Liu W, Zhang LK, Peng K. 2020. SFTSV Infection Induces BAK/BAX-Dependent Mitochondrial DNA Release to Trigger NLRP3 Inflammasome Activation. Cell Rep 30:4370–4385 e7.

39. Yu S, Zhang Q, Su L, He J, Shi W, Yan H, Mao H, Sun Y, Cheng D, Wang X, Zhang Y, Fang L. 2023. Dabie bandavirus infection induces macrophagic pyroptosis and this process is attenuated by platelets. PLoS Negl Trop Dis 17:e0011488.

40. Okazaki T, Higuchi M, Takeda K, Iwatsuki-Horimoto K, Kiso M, Miyagishi M, Yanai H, Kato A, Yoneyama M, Fujita T, Taniguchi T, Kawaoka Y, Ichijo H, Gotoh Y. 2015. The ASK family kinases differentially mediate induction of type I interferon and apoptosis during the antiviral response. Sci Signal 8:ra78.

41. Zhang L, Fu Y, Zhang R, Guan Y, Jiang N, Zheng N, Wu Z. 2021. Nonstructural Protein NSs Hampers Cellular Antiviral Response through LSm14A during Severe Fever with Thrombocytopenia Syndrome Virus Infection. J Immunol 207:590–601.

42. Zhang S, Zheng B, Wang T, Li A, Wan J, Qu J, Li CH, Li D, Liang M. 2017. NSs protein of severe fever with thrombocytopenia syndrome virus suppresses interferon production through different mechanism than Rift Valley fever virus. Acta Virol 61:289–298.

43. Yoshikawa R, Kawakami M, Yasuda J. 2023. The NSs protein of severe fever with thrombocytopenia syndrome virus differentially inhibits the type 1 interferon response among animal species. J Biol Chem 299:104819.

44. Wu X, Qi X, Qu B, Zhang Z, Liang M, Li C, Cardona CJ, Li D, Xing Z. 2014. Evasion of antiviral immunity through sequestering of TBK1/IKKepsilon/IRF3 into viral inclusion bodies. J Virol 88:3067–76.

45. Wuerth JD, Weber F. 2016. Phleboviruses and the Type I Interferon Response. Viruses 8.

